# Donor monocyte-derived macrophages promote human acute graft versus host disease

**DOI:** 10.1101/787036

**Authors:** Laura Jardine, Urszula Cytlak, Merry Gunawan, Gary Reynolds, Kile Green, Xiao-Nong Wang, Sarah Pagan, Maharani Pradnya, Christopher A Lamb, Anna K Long, Erin Hurst, Smeera Nair, Graham Jackson, Amy Publicover, Venetia Bigley, Muzlifah Haniffa, A. John Simpson, Matthew Collin

**Affiliations:** Human Dendritic Cell Laboratory, Institute of Cellular Medicine, Newcastle University, Newcastle upon Tyne, NE2 4HH, UK; Institute of Cellular Medicine, Newcastle University, Newcastle upon Tyne, NE2 4HH, UK; Northern Centre for Bone Marrow Transplantation, Freeman Hospital, Newcastle upon Tyne Hospitals NHS Foundation Trust NE7 7DN; NIHR Newcastle Biomedical Research Centre, Newcastle upon Tyne Hospitals NHS Foundation Trust, NE1 4LP, UK

## Abstract

Myelopoiesis is invariably present, and contributes to pathology, in animal models of graft versus host disease (GVHD). In humans, a rich inflammatory infiltrate bearing macrophage markers has also been described in histological studies. In order to determine the origin, functional properties and role in pathogenesis of these cells, we isolated single cell suspensions from acute cutaneous GVHD and subjected them to genotype, transcriptome and *in vitro* functional analysis. A donor-derived population of CD11c+CD14+ cells was the dominant population of all leukocytes in GVHD. Surface phenotype and nanostring gene expression profiling indicated the closest steady-state counterpart of these cells to be monocyte-derived macrophages. In GVHD, however, there was upregulation of monocyte antigens SIRPα and S100A8/9, and transcripts associated with leukocyte trafficking, pattern recognition, antigen presentation, and co-stimulation. Isolated GVHD macrophages stimulated greater proliferation and activation of allogeneic T cells, and secreted higher levels of inflammatory cytokines than their steady-state counterparts. In HLA-matched mixed leukocyte reactions, we also observed differentiation of activated macrophages with a similar phenotype. These exhibited cytopathicity to a cell line and mediated pathological damage to skin explants, independently of T cells. Together, these results define the origin, functional properties and potential pathogenic roles of human GVHD macrophages.

## Introduction

Acute graft versus host disease (GVHD) affects up to 50% of patients receiving allogeneic bone marrow transplantation (BMT) and remains a leading cause of morbidity and mortality (1, 2). GVHD most commonly affects the skin, gut, liver and may also contribute to idiopathic pneumonia syndrome (3). In animal models, donor T lymphocytes play an essential role in immune-mediated damage to host epithelium (4). In human GVHD, mononuclear infiltrates have been observed that include CD8+ T cells, CD4+ Th1, Th2, Th17 and Tregs although no pathognomic effector subset has been observed in all patient cohorts (5–8). Despite the obvious importance of effector T cells, they may not be sufficient to mediate GVHD pathology (4). In almost all GVHD models, pathology occurs in the presence of neutrophils, monocytes and other myeloid components that may infiltrate tissues and amplify local immune responses (9).

Animal models previously demonstrated that immunocompetent donor myeloid cells enhance GVHD, without specifying a particular cell type (10, 11). Macrophages have been implicated through observations that GVHD may be modulated by manipulation of the CSF-1 axis, although opposing effects have been reported, depending on the timing of interventions (12–15). Glucocorticoids also appear to reduce GVHD at least partly through attenuation of macrophage responses (16) and in humanized mice, donor monocytes or DC are absolutely required for xeno-GVHD (17). Knock-out of the ATP receptor P2Y2 on recipient monocytes reduces GVHD lethality (18). Most recently, a specific role of T cell-derived GM-CSF was described in promoting the differentiation of effector macrophages (19).

In humans, a number of reports highlight an increase in myeloid cells bearing macrophage markers, showing that the level of infiltration correlates with clinical severity and outcome (7, 20, 21). However, as shown by high resolution analysis, the myeloid cell compartment of human skin is highly complex with discrete populations of classical dendritic cells (DC), monocyte-derived cells and resident macrophages, (22–26). These observations suggest that the nature of myeloid infiltrates cannot be adequately resolved using in situ microscopy; hence their origin and immune functions in GVHD remain undefined.

The role of (recipient) myeloid cells in responding to danger signals is integral to most models of GVHD, but is not known whether human GVHD infiltrates bearing macrophage markers are recipient, donor, immunogenic or anti-inflammatory. Although donor myelopoiesis usually dominates the peripheral blood compartment during GVHD, recipient dermal macrophages have very slow kinetics of turnover in humans (22) and potentially expand during inflammation (27). Macrophages are capable of mediating a wide spectrum of tolerogenic or pathogenic responses (28). By extrapolation from mouse models, macrophages are likely to promote GVHD. However, their ability to stimulate local effector T cells and mediate direct epithelial damage remain untested in humans.

Here we employ direct methods of isolation and testing to show that acute GVHD lesions in human skin are dominated by CD11c+CD14+ myeloid cells with the genotype, phenotype and transcriptional profile of donor monocyte-derived macrophages. These cells have potent immunological functions that are likely to contribute to the pathogenesis of GVHD and may offer opportunities for therapeutic intervention.

## Results

In order to investigate the properties of myeloid cells in human cutaneous acute GVHD, mononuclear cells of the human dermis were defined by immunohistochemistry, immunofluorescence microscopy and flow cytometry in healthy controls, BMT recipients without acute GVHD, and BMT recipients with GVHD (**Table S1, S2**). BMT control patients without GVHD were biopsied on median day 83 post-transplantation (range 28-148 days). GVHD skin biopsies were taken at the onset of an acute onset erythematous rash, prior to initiation of therapy. Classical acute GVHD, immunosuppression withdrawal acute GVHD, and acute GVHD following donor lymphocyte infusion were all included. A pathological diagnosis of acute-type GVHD was confirmed by standard histological criteria in all cases and patients with clinical or histological features of chronic GVHD were excluded. The median day of biopsy was day 53 (range 13-304; Mann-Whitney p = 0.27 compared with BMT controls). *In situ* analysis showed an increase in CD3+ T cells and CD11c+ myeloid cells in a perivascular and epidermal interface distribution in GVHD (Figure 1a). The nature of the leukocytic infiltrate was also documented using 4-colour immunofluorescence of whole-mount specimens. There was marked infiltration of perivascular spaces by CD11c+ cells that usually remained distinct from FXIIIA-expressing resident macrophages (22) (Figure 1b). Further comparison of CD11c, FXIIIA and CD163 antigen expression by this approach is shown in **Figure S1a-c**.

**Fig. 1.**
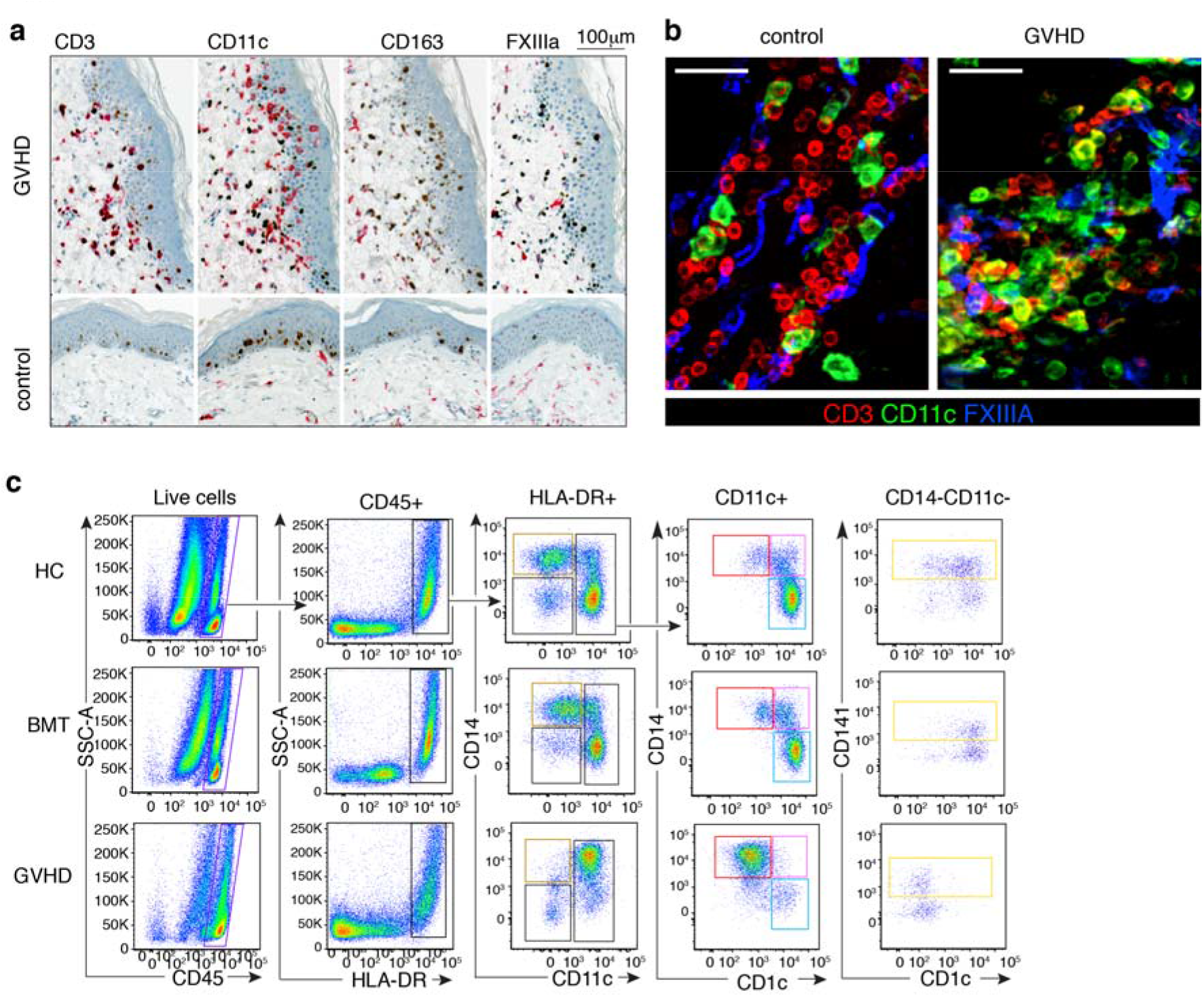
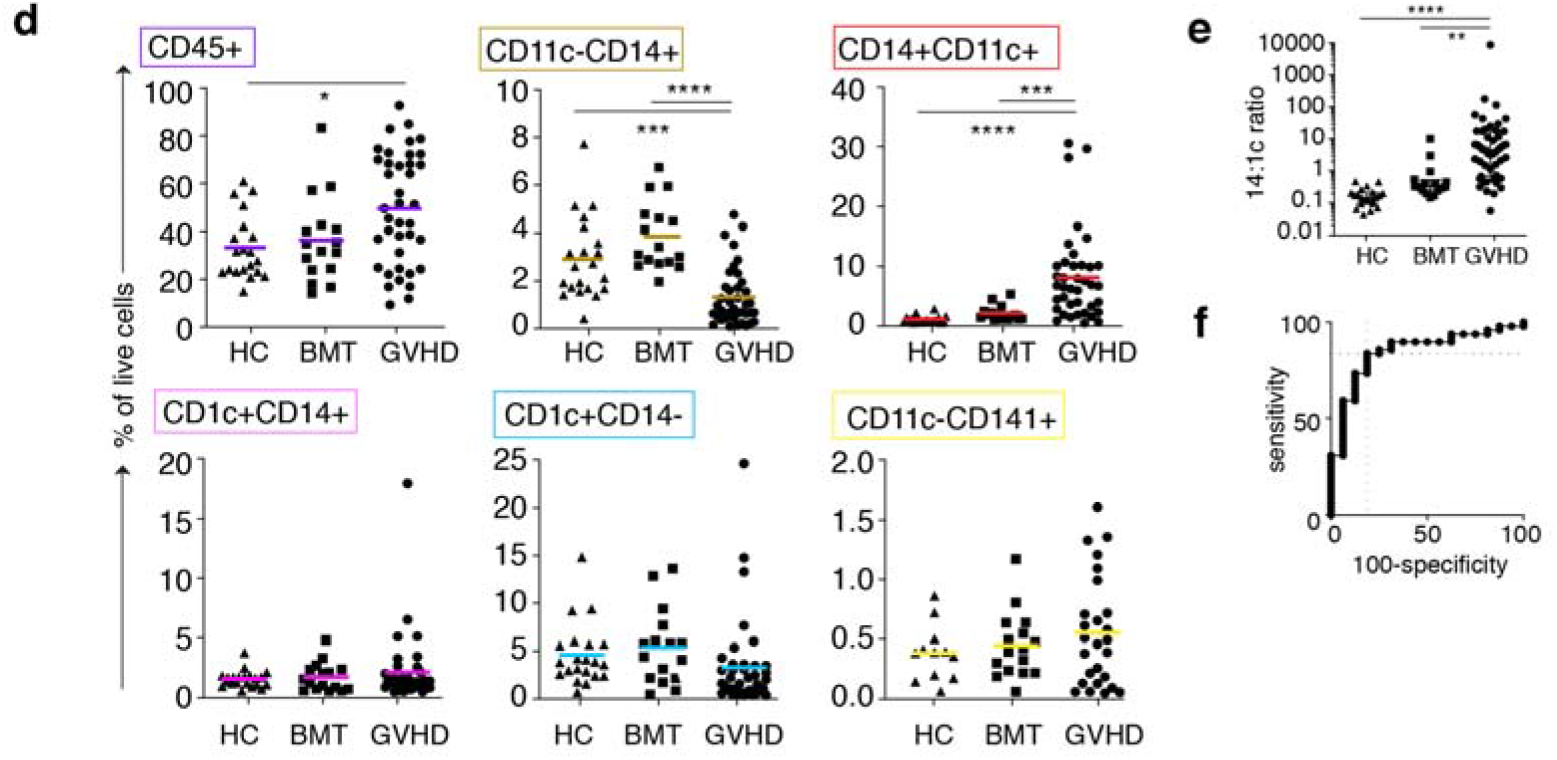
Mononuclear infiltrates in GVHD contain abundant CD14+CD11c+ myeloid cells. Microscopic and flow cytometric evaluation of cutaneous GVHD lesions. Abbreviations: HC: healthy controls; BMT: patients without GVHD; GVHD: patients with GVHD **a** Acute GVHD (top row) and healthy control skin (bottom row). Immunohistochemistry with antibodies to CD3, CD11c, CD163 and factor XIIIa (red chromagen) co-stained with antibody to Ki67 (brown chromagen). **b** Whole mount immunofluorescence of dermis from HC (i) and GVHD (ii) with antibodies to CD3 (red) CD11c (green) and FXIIIA (Blue) **c** Enzymatically digested dermis analyzed by flow cytometry from GVHD, BMT or HC as indicated. Starting from CD45+ mononuclear cells (purple gate), HLA-DR+ cells were gated as shown to arrive at CD11c-CD14+ resident macrophages (brown), CD11c+CD14+CD1c- monocyte-macrophages (red), CD11c+CD14+CD1c+ double positive cells (pink), CD1c+CD14-cDC2 (cyan) and CD141+ cDC1 (yellow; from the CD14-CD11c-gate). Representative samples of more than 60 experiments are shown. **d** Quantification of dermal mononuclear cells from patients with GVHD (n=39), BMT (n=16) or HC (n=21) as a percentage of live cells. Mean and SEM for each group is shown. Groups were compared by one-way ANOVA and p-values from Tukey’s multiple comparisons tests are shown: * p<0.05, **p<0.01, ***p<0.001, ****p<0.0001. **e** Ratio of CD11c+CD14+ cells to CD1c+CD14-cells in GVHD, BMT control or healthy control dermis (14:1c ratio). Median and interquartile range for each group is shown. Groups were compared by non-parametric one-way ANOVA and p-values from Dunn’s multiple comparisons test are shown. **f.** ROC curve analysis of 14:1c ratio in GVHD versus BMT controls. Area under the curve=0.85. Maximal sensitivity and specificity at occurred at a ratio of >0.55.

The infiltrates of acute GVHD infiltrate were characterized by flow cytometry of single cell suspensions. Gating on live singlets expressing CD45 and HLA-DR revealed SSC low lymphocytes and HLA-DR+ SSC high myeloid cells as previously described (22, 25). Surprisingly, the proportion of cells falling in the lymphoid gate was not significantly increased in GVHD relative to BMT controls or healthy donors (**Figure S1e**) although a relative expansion of IFNγ-secreting CD4+ T cells was observed in relative to both controls **(Figure S1d-f).** Myeloid cells were further divided on the bivariate plot of CD14 versus CD11c allowing identification of subsets previously described in healthy control skin, without relying upon autofluorescence to capture resident macrophages (22–24, 26). The linkage between this new gating strategy and previously identified myeloid cell populations is explained in **Figure S2a-d**.

In contrast to the modest changes in overall cellularity and T cell populations, CD11c+CD14+ myeloid cells were expanded more than 10-fold compared with healthy control skin or BMT skin without GVHD (Figure 1c, d**, red gates; Figure S1a-c).** This GVHD-related subset lacked CD1c expression and mapped to parameter space containing monocyte-macrophages in the steady-state (25). Cells in the CD14+ CD11c-gate contained FXIIIA+ CD163+ macrophages with high melanin content and auto-fluorescence, representing ‘fixed’ or resident macrophages (22, 29, 30). These were relatively depleted in GVHD as were classical DC2 (cDC2; CD11c+ CD1c+ CD14-) and cDC1 (CD141+ cells in the CD14-CD11c-gate; Figure 1c, d). A ratio of >0.55 between CD11c+ CD14+ GVHD cells and CD1c+ cDC2 was 84% sensitive and 81% specific for the histological diagnosis of GVHD in skin biopsies post BMT (Figure 1f). Sequential biopsies showed resolution of the GVHD infiltrate in parallel with clinical improvement (**Figure S2e, f**).

We sought to characterize the excess of CD11c+CD14+ cells observed in GVHD further, showing by morphology that they were small macrophages with eccentric dense nuclei, cytoplasmic vacuoles and granules, distinct from larger, melanin-rich macrophages isolated from the CD14+ CD11c-gates (Figure 2a). They retained similar migratory capacity in vitro to their steady-state counterparts (25) (Figure 2b, c). CD11c+CD14+ GVHD cells expressed common macrophage antigens (CD163, CD64 CD206 and CD209) but showed upregulation of monocyte-associated antigens (CD172a, S100A8/9, CD16 (Figure 2d).

**Fig. 2.**
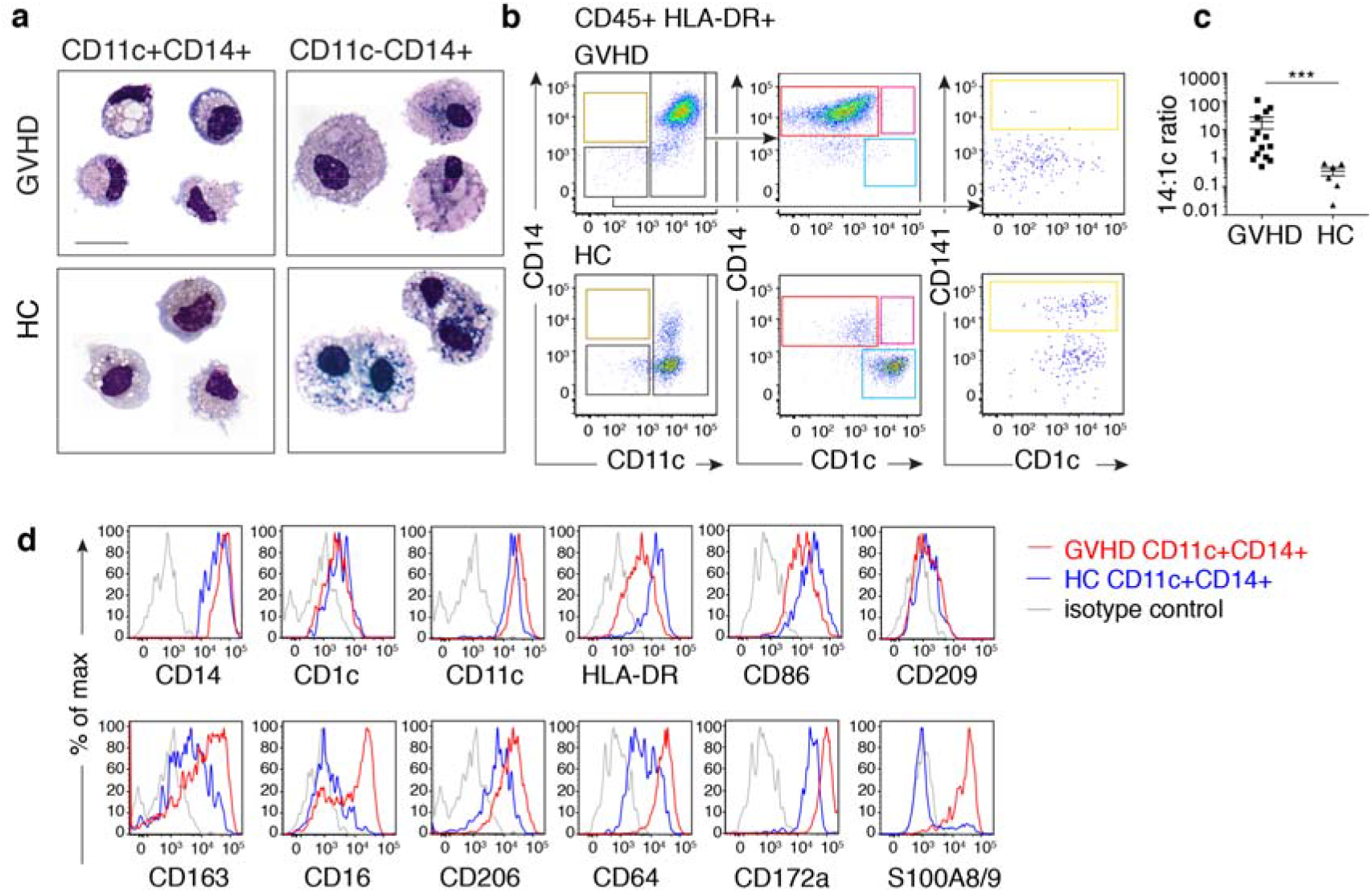
CD14+CD11c+ cells are small migratory macrophages with monocyte antigen expression. **a.** May-Grünwald Giemsa stained cytospins of CD11c+CD14+ and CD11c-CD14+ myeloid cells sorted from GVHD dermis and healthy controls. Representative cells from 2-4 concatenated images are shown. Scale bar represents 20 μm. **b.** Flow cytometry analysis of CD45+HLA-DR+ leukocytes migrating from explanted GVHD or healthy control skin over 48 hours in vitro. Gating as in Figure 1. **c.** Comparison of CD14:CD1c ratio in migrating cells from GVHD skin (n=14) and healthy controls (n=6). Bars show mean ± SEM. *** p=0.0002 by Mann-Whitney test. **d.** Relative expression of selected antigens on CD11c+CD14+ cells migrating from GVHD skin (red line) or healthy control (blue line) compared with isotype control (grey line). Representative data from at least three donors are shown.

In order to define the ontogeny of CD11c+CD14+ cells relative to known populations of macrophages and DC, GVHD and steady state populations were sorted, and expression of 609 immunology-related genes was surveyed by Nanostring. By principal component analysis (PCA), CD11c+CD14+ GVHD cells segregated with steady state monocyte-macrophages and resident dermal macrophages, away from DC populations (Figure 3a). Focusing on a previously defined subset of 29 genes that distinguish between DC and monocytes/macrophage lineages (25), CD11c+CD14+ GVHD cells clustered with steady-state monocyte-macrophages and resident macrophages in unsupervised analysis (Figure 3b). Genotype analysis by XY FISH in sex-mismatched transplants showed a median of 98-100% donor origin of CD11+CD14+ GVHD macrophages, equal to the level of blood myeloid chimerism (Figure 3c, d). Based on these results, we conclude that CD11c+CD14+ myeloid cells in GVHD are donor monocyte-derived macrophages. Recipient T cells were present in the dermis in 3 out of 4 patients at the time of GVHD, in keeping with the recent study of Divito et al., submitted to JCI.

**Figure 3.**
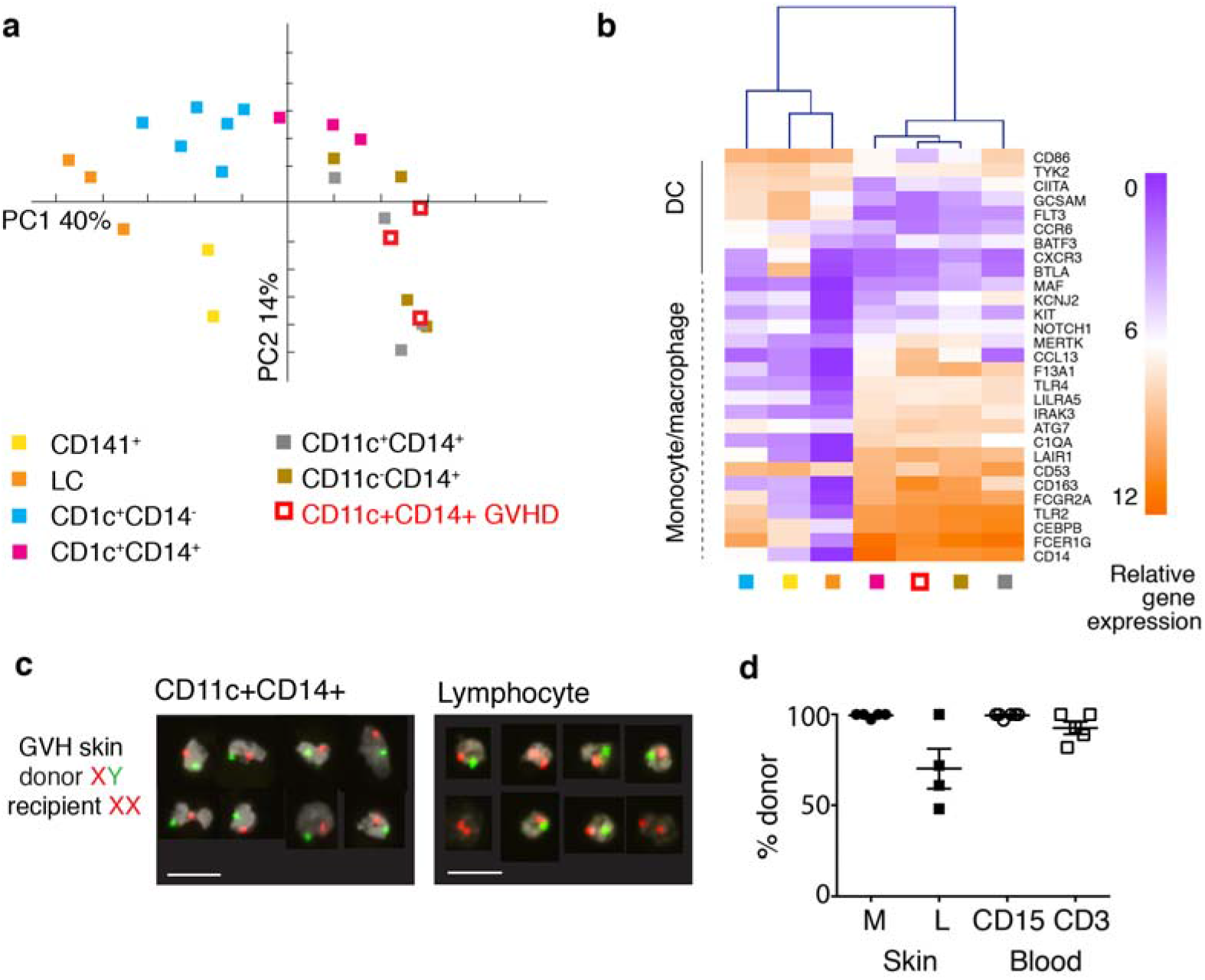
CD14+CD11c+ myeloid cells are donor-derived macrophages. **a.** Principal component analysis (PCA) of immune gene expression by CD11c+CD14+ GVHD cells and 6 myeloid subsets from healthy control skin. Myeloid cells were sorted from healthy control skin as described in Figure 1 and are annotated accordingly. **b.** Heat map showing unsupervised clustering of CD11c+CD14+ cells from GVHD skin and myeloid cells derived from healthy control skin. Mean log2 expression for each subset is shown (n=2 for CD141+, n=3-6 for all other subsets). **c.** Example of FISH showing the XY genotype of GVHD macrophages (CD11c+ CD14+) and lymphocytes sorted from a female recipient transplanted with a male donor. A single field viewed at 10x magnification was concatenated to show 8 representative cells per image. Scale bar represents 20 μm. **d.** Percentage donor origin analyzed by XY FISH of macrophages (M) and lymphocytes (L) sorted from lesional GVHD skin compared with CD15+ myeloid cells (CD15+) and lymphocytes (CD3+) sorted from paired blood samples.

Although myeloid cells have previously been described in human GVHD by histology, their functional properties have not been directly tested. Steady-state CD14+ monocyte-derived macrophages are not potent allo-stimulators compared with dermal CD141+ cDC1 and CD1c+ cDC2 (22, 23). In contrast, GVHD macrophages were capable of stimulating T cell proliferation and expression of activation antigens to the level associated with steady-state DC populations (Figure 4a, b). Gene expression profiling of 2,000-5,000 sorted cells, revealed upregulation of allo-stimulatory functions included antigen presentation (HLA, TAP1), recruitment (CCL24) and stimulation of lymphocytes (CD82), stimulation of pro-inflammatory cytokines (SPP1), and leukocyte extravasation (SELPLG) (Figure 4c). Differential expression of several key chemokines and cytokines was also revealed at the protein level, including IL-6, IL-8, CCL5/RANTES, CXCL10 and IL-10 (Figure 4d).

**Fig 4.**
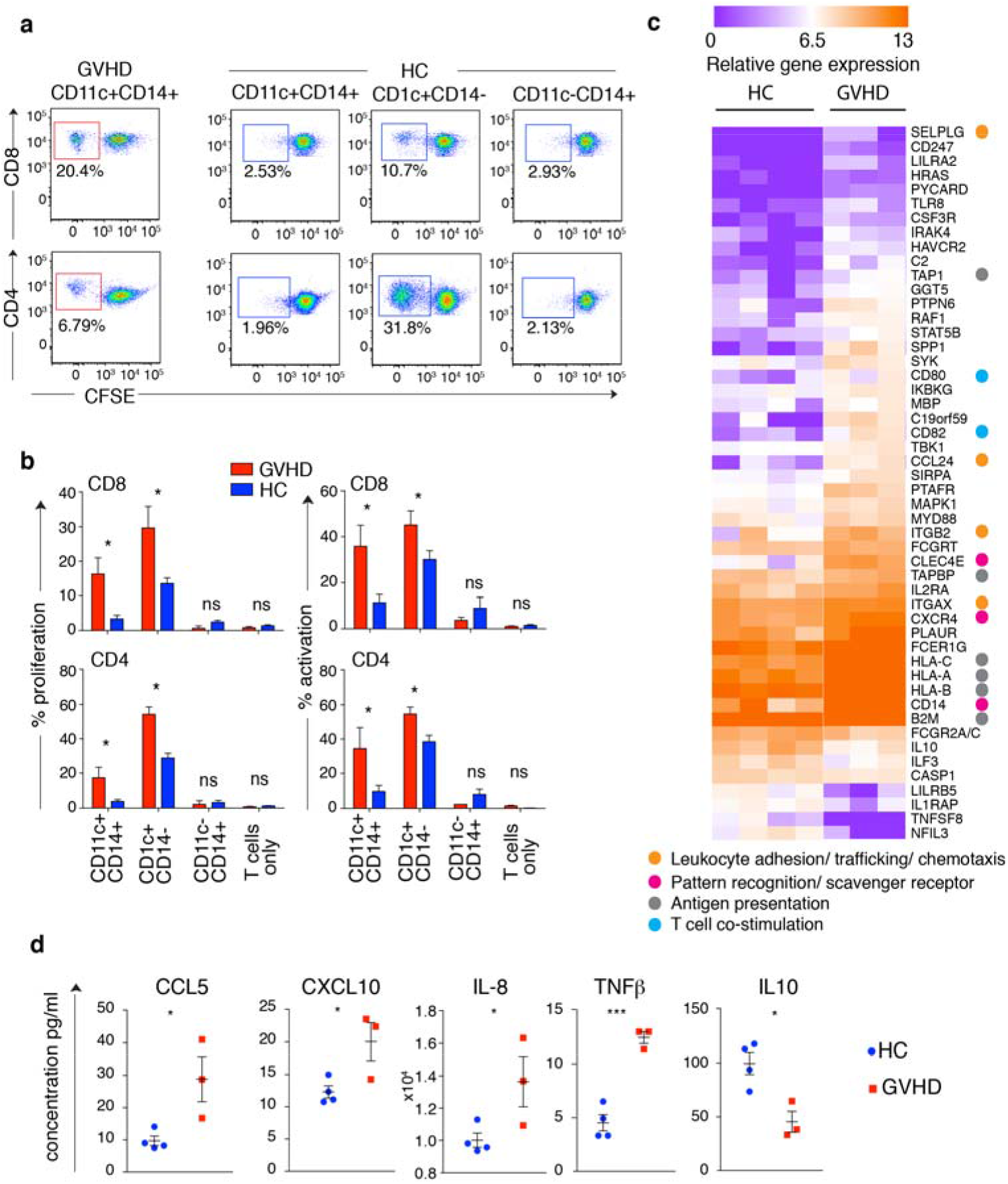
GVHD macrophages activate allogeneic T cells. **a.** proliferation of allogeneic CD4 and CD8 cells estimated by CFSE dilution after co-culture with DC and macrophage subsets sorted from GVHD or healthy controls **b.** summary of T cell proliferation (% CFSE dilution) and activation (% CD69+ CD8+ T cells and % HLA-DR+ CD4+ T cells) from n=3 experiments. *p<0.05 by unpaired t-test. **c.** Heatmap of genes differentially expressed between CD11c+CD14+ monocyte-derived macrophages cells sorted from HC skin (n=4) and GVHD skin (n=3) with fold difference in log2 gene expression >1.3 and p<0.05 by unpaired t-test. Annotations show functional attributes of genes (based on Entrez gene summaries) upregulated in GVHD macrophages **d.** CD11c+CD14+ monocyte-derived macrophages sorted from GVHD (n=3) and healthy control dermis (n=4) were stimulated with LPS in culture over 10 hours. Chemokine and cytokine production was quantified in supernatants by Luminex assay. Bars show mean ± SEM. By unpaired t-test * denotes p<0.05 and *** denotes p<0.001.

The preceding data suggest that GVHD macrophages are derived from blood monocytes and achieve a higher state of functional activation than their steady state counterparts. Further evidence of their likely function in GVHD was sought by deriving allo-stimulated macrophages from monocytes and testing their functional properties. HLA-matched donor and recipient blood was taken prior to transplantation and PBMC stored in order to prepare mixed leukocyte reactions. The cytokine milieu of an HLA-matched MLR was similar to that observed when GVHD skin was cultured (Figure 5a), and a prominent population of macrophages appeared, with similar phenotype to GVHD macrophages (Figure 5b). MLR-activated macrophages also expressed cytotoxic molecules perforin, granzyme A, granulolysin and TRAIL, similarly to GVHD macrophages (Figure 5b-d). Many of these products were already upregulated in CD14+ monocytes isolated from the blood of patients with GVHD, compared with healthy control monocytes (Figure 5d).

**Fig 5:**
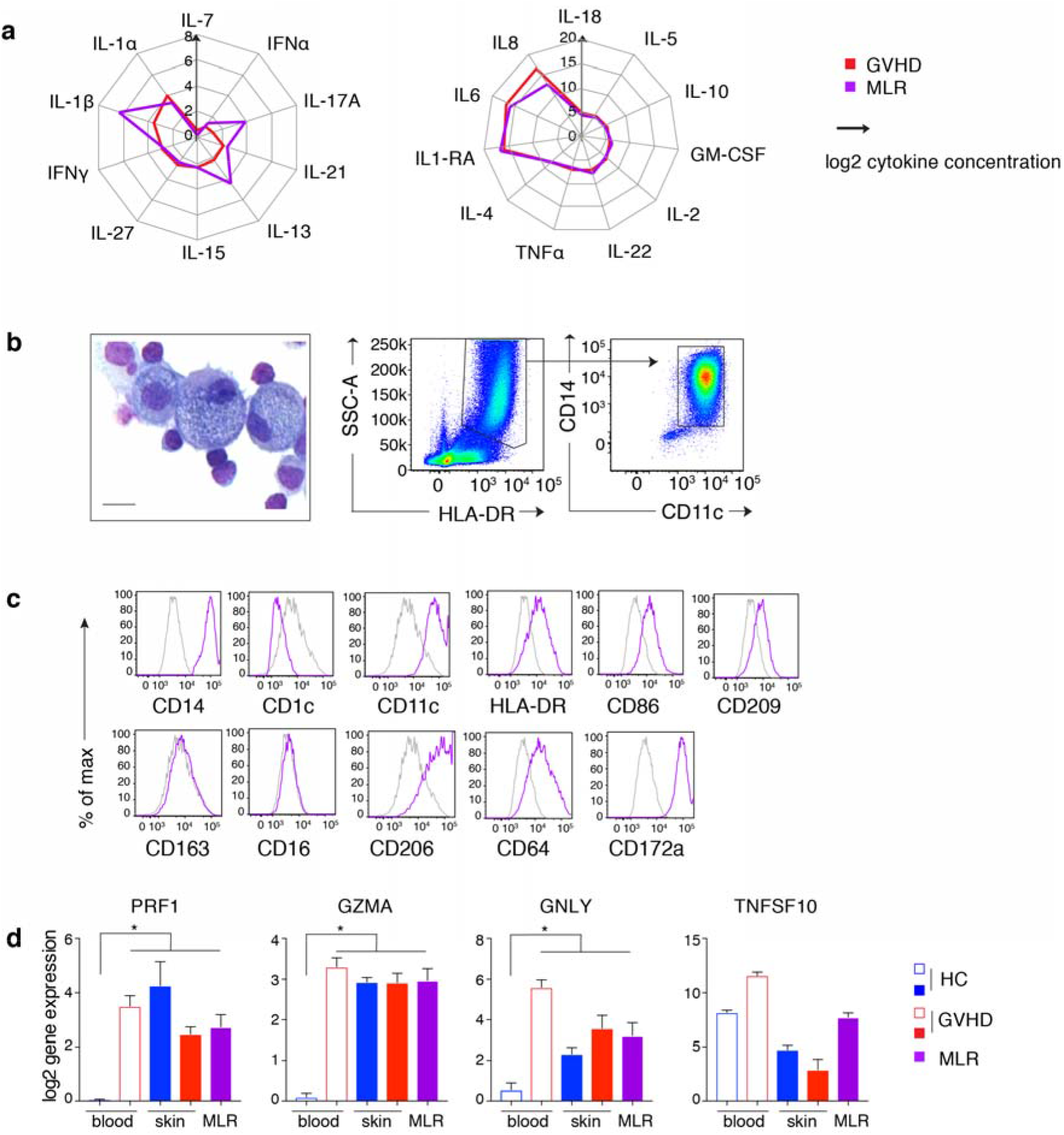
Allo-stimulated monocytes resemble GVHD macrophages. **a.** Radial plots of cytokine quantity in supernatants from GVHD explants cultured for 48 hours (red line) and BMT donor-recipient MLRs cultured for 7 days (purple line). Lines show mean cytokine concentration from n=12 (GVHD) and n=6 (MLR) experiments. IL-9, IL12p70, IL-23, IL-31 and TNF-β, are not shown because they were not detected in any specimens. **b.** May-Grünwald Giemsa cytospin morphology, scatter properties and CD11c/CD14 expression by MLR macrophages, isolated on day 7. Scale bar represents 20 μm. **c.** Expression of selected antigens, previously used to define GVHD macrophages, by allo-stimulated CD11c+CD14+ cells from BMT donor-recipient MLRs (purple line; isotype: gray line) Representative histograms from more than 3 analyses are shown. **d.** Expression of cytotoxic effector genes in CD14+ blood monocytes, skin CD11c+CD14+ cells and MLR macrophages. Columns indicate mean and bars SEM of n=3-6 values; * p <0.05, Mann-Whitney, compared with HC blood.

The expression of cytotoxic molecules prompted us to test the possibility that MLR-activated macrophages might mediate cytotoxicity to epidermal cells. We observed that MLR-activated macrophages were directly cytotoxic to a keratinocyte cell line in vitro, in a dose-dependent manner (Figure 6a, b). In order to test a setting more relevant to GVHD, we adapted the *in vitro* skin explant model. When a small explant of intact skin is exposed to a clone of minor-histocompatibility antigen-specific T cells, GVHD-like epidermal pathology is observed in an HLA-restricted and antigen-specific manner (31). GVHD pathology is also observed, in proportion to HLA-matching and sex differences, when recipient skin is exposed to donor leukocytes pre-sensitized to recipient antigens in a mixed leukocyte reaction (MLR) (32). Although it has been assumed that GVHD pathology in vitro is exclusively mediated by T cells in the MLR, we were surprised to observe nearly equivalent cytopathic effects when the ‘donor’ MLR was sorted into macrophage and T cell components (Figure 6c, d).

**Figure 6.**
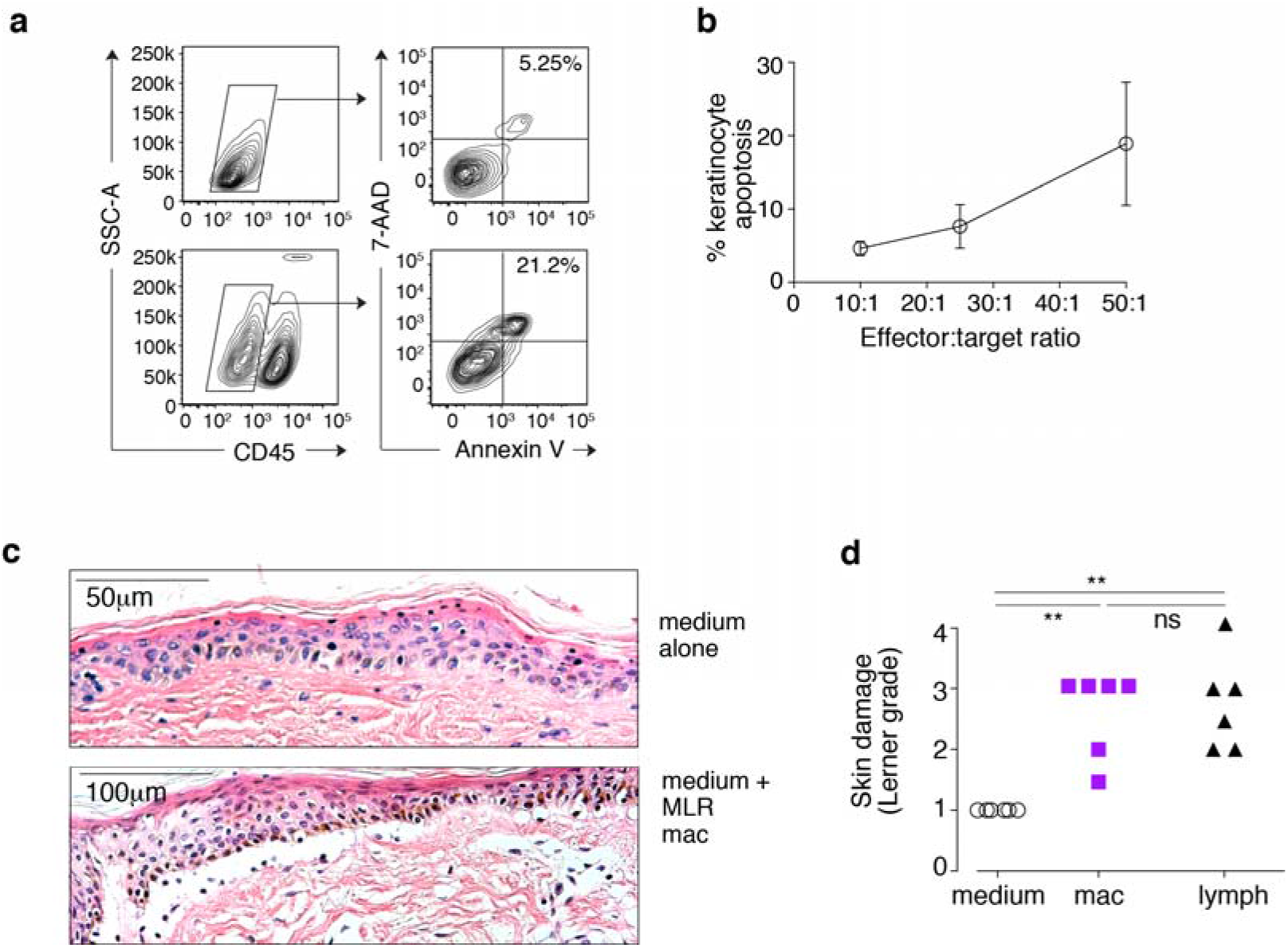
Cytoxicity of allo-activated macrophages in vitro. **a.** Direct cytotoxicity of MLR macrophages to the keratinocyte cell line HaCaT was assessed by co-culture of HaCaT and MLR macrophages at a range of effector to target ratios for 5 hours. Keratinocytes were identified as CD45-cells by flow cytometry and the proportion of apoptotic keratinocytes quantified by Annexin-V and 7-AAD staining. Representative flow cytometry plots from keratinocytes alone (top row) and keratinocytes cultured with MLR macrophages at a 50:1 ratio (bottom row). **b.** Summary of keratinocyte apoptosis across 3 independent experiments showing does-depedent relationship; bars show SEM. **c.** MLR outputs were sorted to yield macrophages and lymphocytes and co-cultured with shave biopsies of recipient skin for 3 days. Explants were fixed and stained with Haematoxylin and eosin. Representative images from explants cultured for 3 days in medium alone or medium + MLR macrophages. **d.** Summary of histological damage to the dermo-epidermal junction graded on the Lerner scale from 6 independent experiments. p values from one-way non-parametric ANOVA and Dunn’s multiple comparison tests are shown.

## Discussion

In this study we have defined the role of myeloid cells in human cutaneous acute GVHD by characterizing mononuclear infiltrates from primary tissue, isolating the dominant myeloid cell, and defining its origin, transcriptional profile and functional properties. The data indicate that human GVHD lesions are highly infiltrated with donor monocyte-derived macrophages with enhanced allo-stimulatory activity and the potential to mediate epidermal pathology.

Myeloid cells found in GVHD have previously been characterized as macrophages based on descriptive histopathology with a small number of surface antigens. These studies lack further details of the biological characteristics or potential pathogenic role of macrophages in GVHD (7, 20–22). Indeed, evidence that macrophages enhance local effector immune functions is surprisingly hard to find in any scenario of inflammation in human tissues. Where they have been isolated from primary human tissues, their function has been described as regulatory, in comparison with DC (33, 34). Our findings that GVHD macrophages have functional attributes capable of promoting GVHD provide an important corroboration of recent mouse models describing the dependence of GVHD pathology upon donor myeloid cells activated by T cell-derived GM-CSF (19).

Numerically, macrophages show the greatest fold increase in GVHD of any mononuclear cell and constitute the most consistent ‘cellular signature’ of acute GVHD relative to recipients without GVHD or healthy control skin. The macrophage:DC ratio is sensitive and specific relative to BMT control skin without GVHD, increasing more than 100-fold in the presence of GVHD. Previous studies of human GVHD have placed emphasis upon the potential existence of a pathognomonic subset of T cells in GVHD, although none has been consistently identified (6–8). Unlike animal models of BMT in which additional splenic T cells are added to initiate GVHD (4), human BMT recipients are typically severely lymphopenic owing to the routine use of post-transplant cytotoxic agents, calcineurin inhibitors, and T cell depletion strategies. Numerically, the T cell infiltrate is surprisingly modest and insignificantly different to healthy human skin, especially in classical early acute GVHD. The observations that MHC class I and II mismatches both increase the risk of GVHD and that CD4 or CD8 selective depletion does not abrogate GVHD are in keeping with multiple pathways of T cell alloreactivity that may vary from patient to patient (4). The striking feature is that all appear to result in the profound recruitment of inflammatory macrophages.

Here, GVHD macrophages were defined as CD11c+CD14+ cells, based on the most direct means of distinguishing the infiltrate from auto-fluorescent CD11c-resident macrophages by flow cytometry. Several lines of evidence point to a monocyte origin; most notably they are donor-derived and therefore unlikely to arise by proliferation of resident recipient macrophages. Additional staining demonstrated high expression of monocyte antigens S100A8/A9 and SIRPA (CD172), consistent with recent emigration of monocytes from the blood (35, 36). Nanostring gene expression analysis confirmed transcription of a core set of macrophage-related genes including MAF, MERTK, F13A1, CD163 and CD14. Although GVHD macrophages have higher expression of monocyte antigens and a number of important functional differences, they are most closely related to CD11c+CD14+ dermal cells found in steady-state tissues and previously reported to have a transient, monocyte origin (25). Enhanced monocyte priming and recruitment to tissues is also suggested by the phenotypic activation of peripheral blood monocytes, previously been reported in patients with active GVHD (37–39). We observed a similar phenomenon in the expression of cytotoxic genes by GVHD monocytes compared with those from healthy controls. In corroboration, we also observed that monocytes differentiating into macrophages in HLA-matched MLRs had a similar phenotype and functional properties to GVHD macrophages.

The data indicate that single surface markers previously used to define GVHD macrophages by histology often have variable expression under more detailed scrutiny. In keeping with previous reports (7, 20–22), CD163 was detectable by flow cytometry, but was less consistent than CD11c+ in identifying GVHD infiltrates by immunohistochemistry. CD163 is expressed by resident macrophages (40) and chronic inflammatory macrophages found in psoriasis (41), coeliac disease (42) and Crohn’s disease (43). Higher CD163 expression was associated with longer intervals post-transplant, consistent with the description of abundant CD163 expression in advanced GVHD lesions (20, 21). These findings are also in keeping with the original characterization of CD163 (clone RM3/1) as a ‘late phase’ macrophage antigen with more delayed kinetics of expression (44).

GVHD macrophages demonstrated enhanced T cell stimulatory functions, compared with steady-state CD11c+CD14+ cells, including greater expression of pattern recognition, leukocyte adhesion and trafficking, antigen presentation and T cell co-stimulation genes, production of chemokines and capacity to stimulate allogeneic T cell proliferation. The potential of macrophages to mediate direct cytotoxic effects is described in classical studies but, until recently, has received little attention in the field of GVHD. Early studies showed that GVHD induced priming of macrophages resulting in direct cytotoxicity following LPS challenge (45) and subsequent work elaborated on the secretory properties of activated macrophages (46). However, contemporary models of GVHD present the contribution of macrophages to GVHD almost exclusively in terms of sensing danger and enhancing accessory cell function (9–11). It was not possible to harvest sufficient GVHD macrophages directly from biopsies to test their effector function so we generated allo-stimulated macrophages from monocytes in HLA-matched MLRs, as a surrogate. MLR-stimulated macrophages were capable mediating direct cytopathicity with a cell line and surprisingly, caused a similar degree of immunopathology as T cells in the skin explant model of GVHD. Although the latter lacks all the complexity of GVHD in vivo, it is the only human system amenable to manipulation. Furthermore, the degree of pathological damage consistently reflects levels of major and minor histocompatibility antigen matching and has been used previously to dissect HLA-restricted antigen-specific GVHD responses (31, 32). This result revises the assumption that T cells are the only relevant effectors when the an MLR product is added to explanted skin.

In classical animal models of GVHD, myelopoiesis is invariable present during the effector phase of GVHD, even when the sole instigator of GVHD is a T cell clone directly targeted to epithelium (47, 48). Investigators have now revealed non-redundant functions for myeloid cells in GVHD pathogenesis (12–15, 17, 19). The conceptual advance that T cells are necessary but may not be sufficient for GVHD has important therapeutic implications. Unlike T cells in the human adult, which may take more than 2 years to recover after transplantation, myeloid cells are continuously generated, and rapid immune reconstitution is possible following myeloid-targeted interventions. The ability to isolate discrete mechanisms that govern the infiltration of tissues by myeloid cells, such as GM-CSF dependence, may also offer a means of minimising GVHD without compromising GVL (19).

Inflammatory macrophages with a similar phenotype to GVHD macrophage have been characterized in other pathological states such as psoriasis (41) coeliac disease, (42) and Crohn’s disease (43). In coeliac disease, acute gluten challenge induces an increase of CD11c+CD14+ cells with modest expression of CD163, reminiscent of the population found in GVHD (49). These observations suggest that similar inflammatory myeloid cells are found in other conditions. Owing to the distinct clinical context of BMT, these are unlikely to become confused with GVHD, but may present opportunities to target common pathways across a wide range of conditions.

Intriguingly, in the field of solid organ transplantation, macrophages have been reported to sense histocompatibility differences. This mechanism has not been explored in GVHD but could initiate inflammatory responses especially in the context of lymphopenia. (Zecher et al., 2009; Liu et al., 2012; Oberbarnscheidt et al., 2014; Fox et al., 2001). Furthermore, as shown in the recently submitted manuscript of Divito et al., abundant host tissue resident T cells, survive conditioning and may initiate GVHD by stimulating an initial host versus graft (HVG) reaction by recipient T cells. Consistent with this hypothesis, it has been observed that DP mismatches in the HVG direction are associated with higher levels of GVHD than fully DP-matched transplants (Reid et al, manuscript in preparation). The possibility that a tissue-based HVG reaction can cause pathology in distinguishable from GVHD is also suggested by the observation that lethal GVHD may be mediated by a ‘bystander’ MLR effect in which recipient epithelium is not directly recognizable by T cells, in animal models (50).

In summary, the results presented here demonstrate that GVHD lesions contain abundant donor macrophages, likely to be derived from activated circulating classical monocytes. GVHD macrophages secrete chemokines, are capable of stimulating T cells and can mediate direct cytotoxicity. Together, these results shed new light on human macrophage functions and suggest that they may be a universal therapeutic target for the prevention and treatment GVHD.

## Methods

### Human subjects

Sequential patients undergoing allogeneic BMT were recruited from the Northern Centre for Bone Marrow Transplantation at Newcastle upon Tyne Hospitals NHS Foundation Trust over a 3 year period between 2013 and 2016. 5-15mm^2^ skin shaves were obtained using 1% lidocaine local anesthesia and a Dermablade^TM^ from BMT recipient controls with no evidence of GVHD and BMT recipients with GVHD. Biopsies were transported in serum-free medium (X-VIVO, Lonza) and analysed within 24 hours. An independent clinical pathologist provided diagnosis and histological grading of GVHD in controls and GVHD biopsies. Healthy control skin was obtained from patients undergoing mammoplasty or abdominoplasty as previously described (22). Specimens comparable to clinical biopsies were obtained by immobilizing skin strips on a cork block covered with sterile silicon and performing skin shave biopsies.

### Cell lines and mononuclear cells

Unless stated otherwise all cells were cultured in RPMI 1640 (Lonza) with 100IU/ml penicillin, 10μ/ml streptomycin, 2mM L-glutamine (Invitrogen), 10% heat-inactivated fetal bovine serum (Sera Lab). MLR macrophages were generated from co-culture of peripheral blood mononuclear cells (PBMCs) from HLA-matched BMT donor and recipient pairs. Recipient PBMCs were irradiated (20Gy) and used as stimulators for donor PBMCs at a 1:1 ratio. Cultures containing 1-5×10^7^ donor PBMCs were maintained in RPMI 1640 with 10% human AB serum (Sigma) for 7 days.

### Dermal cell isolation

Skin shave biopsies were used whole or split into dermis and epidermis by treatment with dispase 0.5-1.0mg/ml in RPMI for 60-90 minutes at 37°C (Gibco) before digestion in medium with 1.6mg/ml type IV collagenase (Worthington) for 12-16 hours at 37°C in RPMI with 10% heat-inactivated fetal calf serum. Gentle dissociation and passage through a 100 μm filter generated single cell suspensions. For some experiments, an enzyme-free preparation of leukocytes was required. Skin was cultured as above but without collagenase. After 48 hours, migratory leukocytes were harvested from the supernatant. Supernatants were stored at −80°C and later used for cytokine analysis by Luminex assay (see below)

### Immunohistochemistry

Formalin-fixed and paraffin-embedded skin shave biopsies from GVHD diagnostic material were used. 4 μm sections were made. Antigen retrieval and staining were performed using the BenchMark autostainer (Ventana). CD3, CD163, CD11c, factor XIII and Ki67 primary antibodies and the ‘ultraview’ detection kit were used (X).

### Fluorescence microscopy

200μm sheets of whole skin were fixed in 2% paraformaldehyde (X) and 30% sucrose (X) in PBS (X) for 12-18 hours at 4**°**C, washed in PBS and stored at 4**°**C until staining. Specimens were blocked in 0.5% bovine serum albumin (X) with 0.3% TritonX-100 (X) in PBS and stained at 4**°**C for 12-18 hours with the following primary:secondary combinations: CD11c-biotin (X):Streptavidin Cy5 (X); Factor XIII: donkey anti-sheep Alexa Fluor 647 (X); CD3 (X): donkey anti-mouse Alexa Fluor488 (X). Sections were immersed in DAPI-containing mounting medium for 12-18 hours at 4**°**C, then visualized on a Zeiss Axioimager Z2 using the Apotome function and Axiovision version 4.8 software.

### Flow cytometry analysis and sorting

Antibodies used for flow cytometry are listed in **Table S3**. DAPI was added for dead cell discrimination (X). Flow cytometry analysis and sorting was performed using BD FACS Canto II, BD Fortessa X20, BD FACS Aria II and BD FACS Fusion running FACSDiva version 7 and analyzed with FlowJo (TreeStar version X).

### X/Y fluorescence in situ hybridization

Sorted cells were spun onto slides, fixed in methanol and acetic acid and prepared with dual labeled XY FISH probes on a ThermoBrite system (Abbott) in accordance with manufacturer’s instructions.

### Gene expression by NanoString

Gene expression was quantified by NanoString, using the Human Immunology v2 panel with a custom 30-gene add-on targeting monocyte/macrophage and DC genes (ASIP, C19orf59, CCL17, CD1C, CD207, CLEC10A, CLEC9A, CLNK, COBLL1, CXCL5, DBN1, F13A1, FGD6, FLT3, GCSAM, GGT5, Ki67, LPAR2, LYVE1, MAFF, MERTK, NDRG2, PACSIN1, PPM1N, PRAM1, S100A12, SIRPA, TMEM14A, UPK3A and ZBTB46). 3,000-10,000 sorted cells were pelleted and lysed in 5μl RLT buffer (Qiagen) + 1% beta-mercaptoethanol yielding 50-150ng total RNA for analysis. Hybridization was performed according to manufacturer’s instructions using a NanoString Prep Station and Digital Analyzer. nSolver Analysis software version 3.0 was used for background correction and normalization.

### Lymphocyte proliferation assays

Healthy volunteer T cells were prepared from healthy blood donors by immunodensity negative selection (Human T cell enrichment cocktail, Stemcell 15021) and labeled with 1μM CSFE (Invitrogen). T cells were co-cultured with sorted macrophages or DCs at a 25:1 ratio for 7 days, incubated with antibodies to CD3, CD4, CD8, HLA-DR and CD69 (**Table S3**), and analyzed by flow cytometry.

### Cytokine and chemokine quantification

Quantification was performed on medium from isolated macrophage populations stimulated ex-vivo and medium conditioned by explanted GVHD skin and by donor-recipient MLR. For macrophage stimulation, FACS sorted skin macrophages were stimulated with 100ng/ml LPS from E. coli (Sigma) and harvested at 10 hours. Supernatants were cryopreserved at −80**°**C and batch-analyzed by Luminex assay (ProcartaPlexTM 34-plex, EBioscience) using a Qiagen Liquichip 200 running Luminex 100 integrated system software version 2.3. Procartaplex Analyst version 1.0 was used to define standard curves.

### Skin explant assay

Blood from BMT donor and recipient pairs was prospectively collected into EDTA, prior to transplnatation. Recipient PBMCs were irradiated (20Gy) and used as stimulators for donor PBMCs at a 1:1 ratio. Cultures containing 1-5×10^7^ donor PBMCs were maintained in RPMI with 10% human AB serum (Sigma) for 7 days. Macrophages and T cells were sorted from MLR outputs as HLA-DR+CD14+CD11c+ cells and SSC^lo^CD3+ cells respectively. Shave biopsies of recipient skin were taken prior to BMT conditioning. Biopsies were divided and incubated with medium alone (negative control), 2×10^5^ MLR lymphocytes or 2×10^5^ MLR macrophages for 3 days in RPMI 1640 with 20% heat inactivated autologous serum. Explants were fixed in 10% buffered formalin, sectioned and stained with haematoxylin and eosin. Severity of histological damage was graded using the Lerner scale by an experienced assessor blinded to experimental conditions (XN Wang).

### Quantification and statistical analysis

FlowJo version 9.6.7 was used for analysis of flow cytometry data. Principal component analysis (PCA), hierarchical clustering and unpaired t-tests were performed in MultiExperiment Viewer software version 4.8 (Saeed et al., 2003). Other statistical analyses were performed in GraphPad Prism, version 7.0. Definitions of center and dispersion, definitions of significance and statistical methods used are provided in figure legends.

### Study approval

All human samples were obtained with informed consent according to the protocols: Improving Haematopoietic Stem cell Transplantation Outcome, Newcastle and North Tyneside Research Ethics Committee 2 reference 14/NE/1136; or Newcastle Biobank, Newcastle and North Tyneside Research Ethics Committee 1 reference 17/NE/0361.

## Author contributions

Conceptualization: LJ, MC, AJS, MH

Methodology: LJ, MC, VB, SP, MH

Investigation: LJ, UC, MG, KG, GR, SP, MP, CAL, AL, VB

Formal Analysis: LJ, XW

Writing- Original Draft: MC, LJ

Writing- Review & Editing MC, LJ

Funding Acquisition: LJ, MC, AJS

Resources: MC, EH, SN, GJ, VB, GR, MH

Supervision: MC, AJS, MH

## Acknowledgements

We acknowledge the Newcastle University Flow Cytometry Core Facility (FCCF) for assistance with the generation of Flow Cytometry data. This work was funded by Wellcome Trust WT097941 (LJ) and Wellcome Trust 101155/Z/13/Z (V.B., U.C.) with additional contribution from the Newcastle upon Tyne Hospitals Healthcare Charity and the NIHR Newcastle Biomedical Research Centre (AJS) and Bright Red (MC).

---

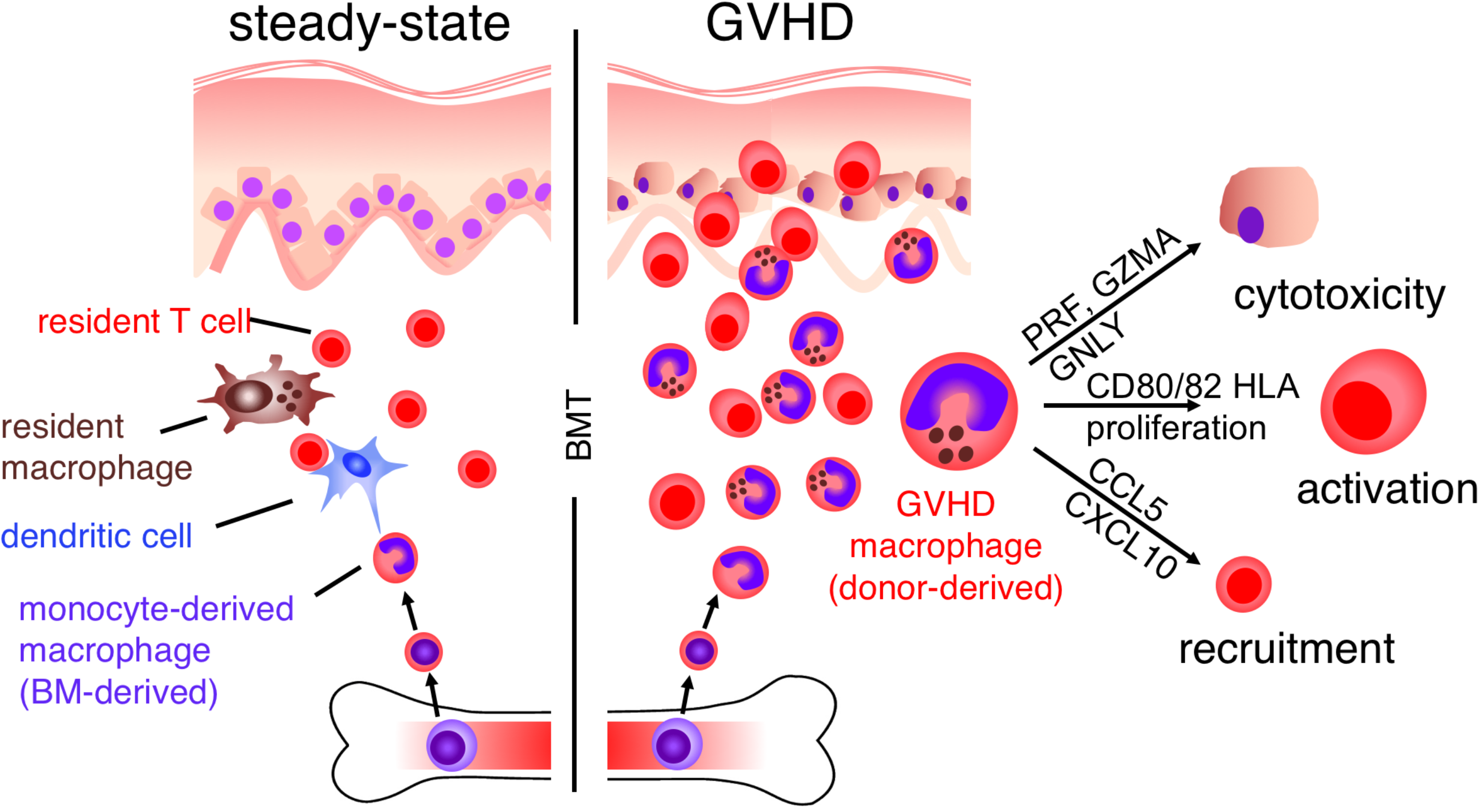

## Notes

***Conflict of interest*:** The authors have declared that no conflict of interest exists

