## Supplementary Table and Figure for "Donor monocyte-derived macrophages promote human acute graft versus host disease"

Supplementary Materials

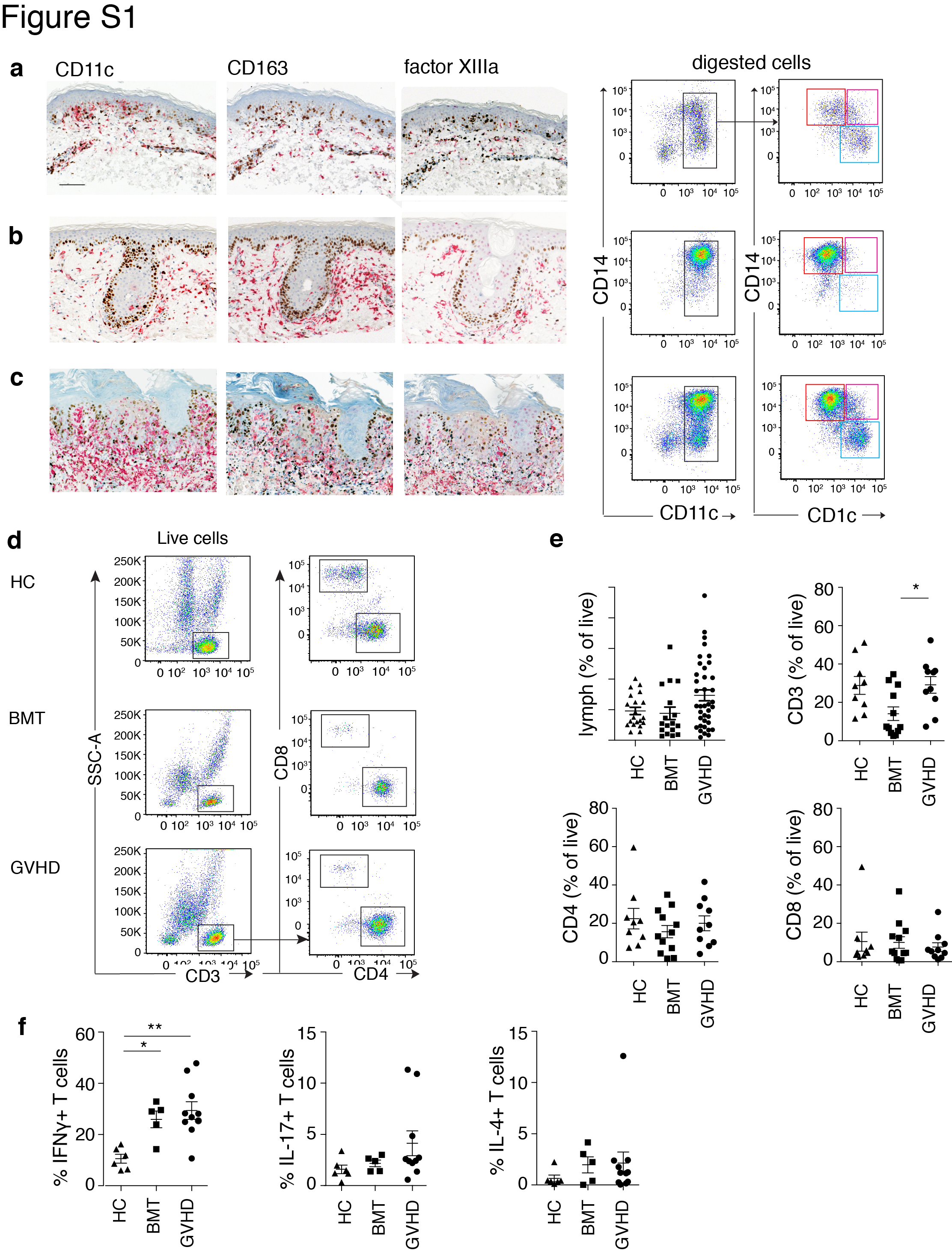

**Fig. S1. Additional characterization of the GVHD infiltrate.**

**Fig. S1. Additional characterization of the GVHD infiltrate.**

**a-c**. Paired IHC and flow cytometry analysis of acute GVHD presenting at different intervals post-transplantation showing increasing burden of macrophages with consistent flow phenotypes. GVHD biopsies were immunostained with antibodies to CD11c and CD163 and factor XIIIa as indicated (red) and co-stained with antibodies to Ki67 (brown). Note epidermal stress indicated by Ki-67 staining. Paired flow cytometry analysis was performed as described in Figure 1.

**a.** 56 days post-BMT with lichenoid dermatitis, histological grade II;

**b.** 214 days post-BMT following ciclosporin withdrawal with eczematoid spongiosis and lichenoid dermatitis, histological grade II;

**c.** 278 post-BMT following donor lymphocyte infusion with acute on chronic lichenoid dermatitis, histological grade II.

**d.** Gating strategy for lymphocytes in flow cytometry of single cell suspensions of digested dermis from patients with acute GVHD, BMT controls and healthy controls. Among live cells, CD3+ SSC^lo^ lymphocytes were divided by CD4 and CD8 expression. Representative examples from more than 8 experiments are shown.

**e.** Relative abundance of lymphocytes by flow cytometry of single cell suspensions of digested dermis from healthy controls (HC; n = 9), BMT controls without GVHD (BMT; n = 8), or patients with acute GVHD (GVHD; n = 10). Error bars show mean ± SEM. * p<0.05 by one-way ANOVA with Dunnett’s multiple comparisons test. Lymph = cells falling in the HLA-DR- SSC low lymphoid gate (see Figure 1).

**f.** Cytokine production by lymphocytes from healthy control (HC, n=6), BMT transplant control (BMT, n=5) or patients with GVHD (GVHD, n=10), following ex vivo stimulation by PMA and ionomycin assessed by flow cytometry. Error bars show mean ± SEM. * p<0.05 and ** p<0.01 by one-way ANOVA with Dunnett’s multiple comparisons test.

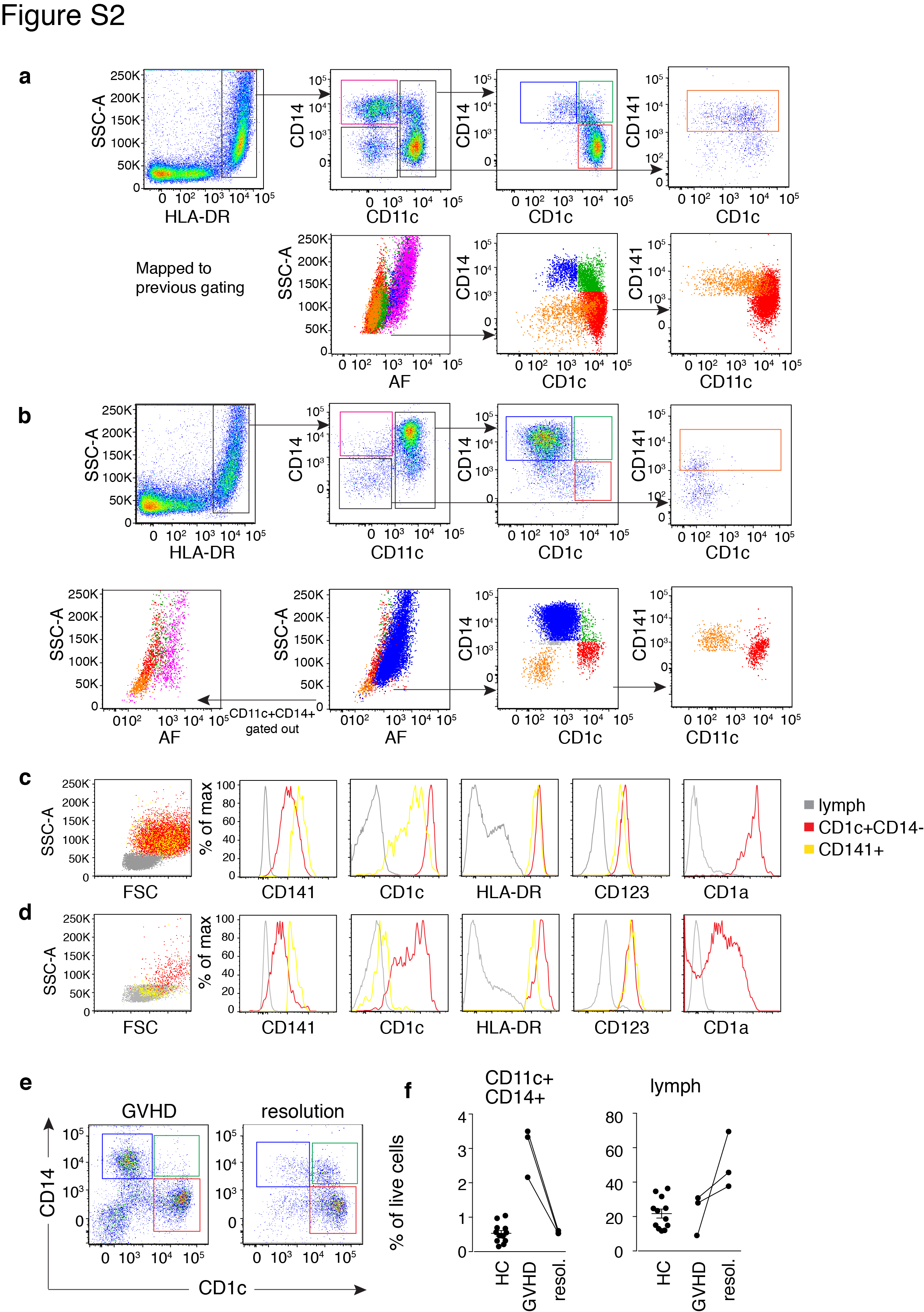

**Fig. S2. Gating strategy for characterization of mononuclear cells in GVHD**

**Fig. S2. Gating strategy for characterization of mononuclear cells in GVHD**

**a,b.** Comparison of previous gating method using autofluorescence to identify dermal macrophages with current schema used to identify myeloid cells in the healthy GVHD dermis.

**a.** Healthy control dermis. Top row shows populations identified with the gating scheme used in Figure 1. Bottom row shows projections of these populations onto the gating scheme used previously, in which autofluorescence was used to identify macrophages (pink) and the small population of monocyte-derived macrophages in normal skin forms a small population (blue) at the interface between macrophages and DC (CD1c+ cDC2: red; CD141+ cDC1: orange). CD14+CD1c double positive cells: green.

**b.** GVHD skin. Top row shows populations identified with the gating scheme used in Figure 1. Bottom row shows projections of these populations onto the gating scheme used previously. Note that the dominant population of monocyte-dervide GVHD macrophages (blue) obscures the distinction between macrophages and DC on the autofluorescence axis. Resident macrophages (pink) can be recovered after the CD11c+ CD14+ GVHD macrophages are gated out (bottom row left-most panel)

c, d Scatter properties and selected antigen expression, confirming the identity of CD1c+ cDC2 (red) and CD141+ cDC1 (yellow) populations

c. Healthy control dermis

d. GVHD dermis

e. Mononuclear cell analysis in paired biopsies at GVHD onset and following resolution of skin inflammation. Dot plots show CD11c+ cells in a representative patient at GVHD presentation and follow-up 86 days later.

f. Proportion of CD11c+ CD14+ GVHD macrophages and lymphocytes at presentation of GVHD and follow-up in 3 paired samples quantified as % of live cells. Accompanying normal ranges in healthy control skin measured by flow cytometry are shown as dot plots with mean ± SEM (n=15).

**Table S1. BMT participant characteristics**

Clinical characteristics of BMT patients with GVHD (‘GVHD’) and controls without GVHD (‘BMT control’) in skin biopsy cohort and BMT recipients undergoing BAL for IPS. Donor type: MRD= matched related donor; MUD= matched unrelated donor. Conditioning: RIC= reduced-intensity conditioning; MAC= myeloablative conditioning; TCD= T cell depleted, TR= T cell replete.

|  | **GVHD**  n=67 | **BMT control**  n=15 |
| --- | --- | --- |
| **Age**; mean (SD) | 53 (12) | 50 (13) |
| **Disease**; n (%)  Acute leukemia  Myelodysplasia  Lymphoproliferative neoplasms  Myeloproliferative neoplasms | 33 (49)  10 (15)  21 (32)  3 (5) | 8 (53)  1 (7)  4 (27)  2 (13) |
| **Donor type**; n (%)  MRD  MUD  Other | 19 (28)  45 (67)  3 (4) | 3 (20)  12 (80)  0 |
| **HLA-match**; n (%)  >10/10  <10/10  Haploidentical | 63 (94)  1 (1)  3 (4) | 15 (100)  0  0 |
| **Conditioning**; n (%)  RIC-TCD  RIC  MAC  Pre-conditioned | 52 (78)  8 (12)  1 (1)  6 (9) | 10 (67)  1 (7)  1 (7)  3 (20) |
| **Serum ciclosporin at biopsy**; μg/L mean (SD) | 132 (151) | 157 (127) |

**Table S2. Conditioning regimens**

| **Group** | **Regimen** |
| --- | --- |
| RIC-TCD  Reduced intensity conditioning, T-depleted | Flu/Mel (Fludarabine 150mg/m^2^, Melphalan 140mg/m^2^, Alemtuzumab 30mg or 60mg, Ciclosporin) |
|  | Flu/Bu (Fludarabine 150mg/m^2^, Busulfan 8-10mg/kg, Alemtuzumab 30mg or 60mg, Ciclosporin) |
|  | Flu/Bu/ATG/MMF (Fludarabine 150mg/m^2^, Busulfan 9.6mg/kg, Fresenius Anti-Thymocyte Globulin 30mg/kg, Mycophenolate Mofetil, Ciclosporin) |
| RIC  Reduced intensity conditioning (T-replete) | Flu/Mel/MTX (Fludarabine 150mg/m^2^, Melphalan 140mg/m^2^, Methotrexate 15mg/m^2^, Ciclosporin) |
|  | Flu/TBI (Fludarabine 90mg/m^2^, 200cGy TBI, Methotrexate, MMF, Ciclosporin) |
|  | RIC HAPLO (Fludarabine 150mg/m^2^, Cyclophosphamide 29mg/kg, 200cGy TBI, post-transplant Cyclophosphamide 100mg/kg, Mycophenolate Mofetil, Tacrolimus) |
| MAC  Myeloablative conditioning | Cy/TBI (12Gy TBI, Cyclophosphamide 120mg/kg, Alemtuzumab 30-60mg, Ciclosporin) |
|  | MAC HAPLO (12Gy TBI, donor lymphocyte infusion, Cyclophosphamide 120mg/kg, Mycophenolate Mofetil, Ciclopsorin) |
| Pre-conditioned | FLAMSA-Bu/ATG/MMF (Fludarabine 120mg/m^2^, Cytarabine 8g/m^2^, Amsacrine 400mg/m^2^, Fludarabine 60mg/m^2^, Busulfan 9.6mg/kg, Genzyme Anti-Thymocyte Globulin 5mg/kg, Mycophenolate Mofetil, Ciclosporin) |
|  | FLAMSA-TBI/ATG/MMF (Fludarabine 120mg/m^2^, Cytarabine 8g/m^2^, Amsacrine 400mg m^2^, 4Gy TBI, Cyclophosphamide 80mg/kg Genzyme Anti-Thymocyte Globulin 5mg/kg, Mycophenolate Mofetil, Ciclosporin) |

**Table S2. Antibodies**

| REAGENT or RESOURCE | SOURCE | IDENTIFIER |
| --- | --- | --- |
| **Antibodies** |  |  |
| CD45 APCCy7 | BD | 557833 |
| CD3 PERCPCy5.5 | BD | 332771 |
| CD4 BV421 | BioLegend | 300531 |
| CD8 PE | BD | 345773 |
| CD127 APC | BioLegend | 351315 |
| CD25 FITC | BD | 345796 |
| CD45 V500 | BD | 560777 |
| CD45 Alexa Fluor 700 | BioLegend | 304024 |
| HLA-DR V500 | BD | 561224 |
| HLA-DR Alexa Fluor 700 | BD | 560743 |
| HLA-DR V450 | BD | 642276 |
| CD14 BV650 | BD | 301835 |
| CD1c PECy7 | BioLegend | 331506 |
| CD11c APCCy7 | BioLegend | 337218 |
| CD11c V450 | BD | 560369 |
| CD123 PERCPCy5.5 | BD | 558714 |
| CD141 PE | BD | 559781 |
| CD141 APC | Miltenyi | 130-090-907 |
| CD3 FITC | BD | 345763 |
| CD4 PE | BD | 555347 |
| CD16 PE-Dazzle594 | BioLegend | 302054 |
| CD14 APCCy7 | BD | 557831 |
| CD14 FITC | BD | 555497 |
| CD1c APC | BD | 559775 |
| CD11c APC | BD | 559877 |
| HLA-DR FITC | BD | 556643 |
| CD86 PE | BD | 555658 |
| CD209 FITC | BD | 551264 |
| CD163 APC | R&D | FAB16078 |
| CD16 FITC | BD | 335035 |
| CD206 APC | BD | 550889 |
| CD64 PE | BioLegend | 305007 |
| CD172a PE | BioLegend | 323805 |
| CD3 V500 | BD | 561416 |
| CD4 PE | BD | 55347 |
| CD8 APCCy7 | BD | 557834 |
| HLA-DR PERCPCy5.5 | BD | 339126 |
| CD69 PE-Cy7 | BD | 557745 |
| CD3 FITC | BD | 345763 |
| CD19 FITC | BD | 345776 |
| CD20 FITC | BD | 345792 |
| CD56 FITC | BD | 345811 |
| HLA-DR V500 | BD | 561224 |
| CD14 Qdot655 | Invitrogen | Q10056 |
| CD16 APCH7 | BD | 560195 |
| CD45 FITC | BD | 345808 |
| Annexin V PE | BD | 56-65875X |
| 7-AAD | BD | 56-66121E |
| S100A8/9 PE | Novus | NBP2-2526PE |
| CD3 unconjugated | BD | 555330; UCHT1 |
| CD11c unconjugated | BD | 555391; B-ly6 |
| Factor XIIIa unconjugated | Enzyme Research | SAF13A-AP (polyclonal) |
| Donkey anti-mouse Dy488 | Jackson ImmunoResearch |  |
| Donkey anti-sheep Alexa Fluor 647 | Invitrogen |  |
| Streptavidin Cy5 | Jackson Immunoresearch |  |
